# Plastid Genome Assembly Using Long-read Data (ptGAUL)

**DOI:** 10.1101/2022.11.19.517194

**Authors:** Wenbin Zhou, Carolina E. Armijos, Chaehee Lee, Ruisen Lu, Jeremy Wang, Tracey A. Ruhlman, Robert K. Jansen, Alan M. Jones, Corbin D. Jones

## Abstract

Although plastid genome (plastome) structure is highly conserved across most seed plants, investigations during the past two decades revealed several disparately related lineages that experienced substantial rearrangements. Most plastomes contain a large, inverted repeat and two single-copy regions and few dispersed repeats, however the plastomes of some taxa harbor long repeat sequences (>300 bp). These long repeats make it difficult to assemble complete plastomes using short-read data leading to misassemblies and consensus sequences that have spurious rearrangements. Single-molecule, long-read sequencing has the potential to overcome these challenges, yet there is no consensus on the most effective method for accurately assembling plastomes using long-read data. We generated a pipeline, ***p***las***t***id ***G***enome ***A***ssembly ***U***sing ***L***ong-read data (ptGAUL), to address the problem of plastome assembly using long-read data from Oxford Nanopore Technologies (ONT) or Pacific Biosciences platforms. We demonstrated the efficacy of the ptGAUL pipeline using 16 published long-read datasets. We showed that ptGAUL produces accurate and unbiased assemblies. Additionally, we employed ptGAUL to assemble four new *Juncus* (Juncaceae) plastomes using ONT long reads. Our results revealed many long repeats and rearrangements in *Juncus* plastomes compared with basal lineages of Poales.

## 1 INTRODUCTION

Plastid genomes (plastomes) are highly conserved, comprising linear, branched or occasionally circular molecules that usually contain a large, inverted repeat (IR) and large and small single-copy regions (LSC and SSC). Due to their conserved structure and low rate of nucleotide substitution, plastome data has made substantial contributions to phylogenetic studies for many plant groups (Jansen & Ruhlman, 2012; Jiang et al., 2022; Liu et al., 2022; Xia et al., 2022; Xu et al., 2022; Yu et al., 2022). Despite the high level of plastome structural conservation in seed plants, rearrangements, including inversions, expansion and contraction of the IR, and IR loss occurred in unrelated lineages of gymnosperms and angiosperms (Ruhlman & Jansen, 2021). Many of these same lineages experienced substantial gene loss with most of these genes functionally transferred to the nuclear genome or substituted by an alternative, nuclear-encoded gene (Ruhlman & Jansen, 2021). Documented transferred/substituted genes include *accD, infA, rpl22, rpl20, rpl32, rpl23, rps7, rps16, ycf1* and *ycf2*.

Genome assembly methods have improved substantially over the past decade (Twyford & Ness, 2017; Freudenthal et al., 2020). NOVOPlasty (Dierckxsens et al., 2017) and GetOrganelle (Jin et al., 2020) are the two most frequently used pipelines for plastome assembly based on Illumina short reads. However these assemblers, which rely on the De Bruijn graph approach (Compeau et al., 2011), do not always yield accurate assembly results when confronted with long repeat regions in plastomes, particularly when those repeats are longer than kmer sizes. In some cases, these tools generate outputs with multiple contigs/scaffolds or hundreds of possible assembly results. The high number of uncertain paths can sometimes be corrected using Bandage (Wick et al., 2015), a software tool that visualizes the depth of read coverage for each contig/scaffold and orders contigs, but the final arrangement of the contigs is often not well-resolved because the Illumina short reads are insufficient to bridge the repeated sequences and their flanking regions. Using short reads with a typical insert size (300-400 bp) is insufficient to obtain a complete plastome assembly for plant species that have large repeats and may be highly rearranged. So far, few plant systematists have recognized this as an issue likely because most plants possess relatively conservative plastome structures with limited repeated sequences and because their primary interest is the extraction of coding sequences for phylogenetic analyses. Long reads generated by third-generation sequencing methods such as Oxford Nanopore Technologies (ONT) or Pacific Biosciences (Pacbio) platforms may help resolve assembly issues as the longer reads are more likely to span long repeats (Liao et al., 2021).

To date, many tools have been developed to assemble organelle genomes using long-read data and hybrid data (both short- and long-read data), including Organelle_PBA (Soorni et al., 2017), Canu (Koren et al., 2017), Unicycler (Wick et al., 2017), and Flye (Kolmogorov et al., 2019; Syme et al., 2021). However, these pipelines for plastome assembly using long-read or hybrid data have some drawbacks. Organelle_PBA was designed exclusively for PacBio data; the Sprai (Miyamoto et al., 2014) and Celera (Miller et al., 2008) assemblers in Organelle_PBA are no longer maintained limiting its extension to assembly with hybrid datasets. The approach of Syme et al. (2021) requires an extra step to manually filter a subset of raw reads matching the plastome (~250X coverage) and sometimes generates multiple contigs in the assembly result. Canu can generate different results depending on different read coverages (Wang et al., 2018). Unicycler was designed for hybrid data, however it takes an extremely long time to finish as input data is increased. All pipelines can likely assemble the conventional plastome, but are not able to assemble atypical plastome structures accurately.

The angiosperm family Juncaceae contains ~500 species within the seven genera *Juncus* L., *Luzula* DC., *Distichia* Nees and Meyen, *Oxychloe* Philippi, *Patosia* Buchenau, *Marsippospermum* Desv. and *Rostkovia* Desv. (Drábková, 2010). *Juncus* is the largest genus and includes ~300 species (Balslev, 2018) and two major subgenera, *Agathryon* and *Juncus* (Drábková et al., 2006; Drábková, 2010). Although many species of Juncaceae have been included in phylogenetic studies using plastid gene sequences and the internal transcribed spacer region of the nuclear ribosomal repeat (Table S1; Drábková et al., 2006; Drábková, 2010; Brožová et al., 2022), species relationships within *Juncus* remain unresolved. Recently, Brožová et al. (2022) incorporated *rbcL, trnL, trnL-trnF*, and ITS1-5.8-ITS2 region to reorganize *Juncus* into seven distinct genera: *Juncus, Verojuncus, Juncinella, Alpinojuncus, Australojuncus, Boreojuncus*, and *Agathryon*.

Not many plastome structures have been reported for *Juncus* (*s.l*.). To avoid the confusion regarding the species names, we did not adopt these latest genera above in our study because a more comprehensive study with more markers is necessary to justify this reranking. So far, the plastome structures are seldom studied in *Juncus* (*s.l*.). Plastomes of just eight *Juncus* species are publicly available in Genbank (Wu et al., 2021; Lu et al., 2021). For example, the focus of Wu et al. (2021) was phylogenetic relationships in the Poales using shared, plastid protein-coding genes and no information was reported on plastome structure. Lu et al., (2021) assembled the plastome of *Juncus effusus* using Velvet (Zerbino, 2010) and NOVOPlasty (Dierckxsens et al., 2017) with GapFiller (Nadalin et al., 2012) without any confirmation by either long range PCR or long-read data leaving the final structure uncertain. Recently, two more *Juncus* (*J. effusus* and *J. inflexus*) nuclear genomes were assembled by Planta et al. (2022), but no plastomes were reported. Adding more complete plastomes of Juncaceae would allow insight into plastome evolution in the family and help gain more phylogenetic insights within Juncaceae and Poales.

To assist in assembling potentially complex plastomes and to explore structural variation in *Juncus*, we created a pipeline, ***p***las***t***id ***G***enome ***A***ssembly ***U***sing ***L***ong-read data (ptGAUL), which assembles plastomes using raw ONT long-read sequencing data. The aims of the study were: 1) test the reliability of the ptGAUL pipeline using 16 published plastomes; 2) employ the pipeline to assemble plastomes of two *Juncus* species (*J. validus* and *J. roemerianus)* sequenced in our study, and assemble two other species (*J. effusus* and *J. inflexus*) from the reads of Planta et al. (2022); and 3) compare plastome evolution in *Juncus* to selected members of the Poales.

## 2 MATERIALS AND METHODS

### 2.1 *Juncus* sample collection and DNA extraction

Young leaves of *Juncus roemerianus* (voucher number: NCU00441655) and *Juncus validus* (voucher number: NCU00434802) were collected from North Carolina, USA and stored in silica gel. Vouchers were deposited in the herbarium of University of North Carolina at Chapel Hill (NCU). Total genomic DNA extraction of dried leaves was performed using a modified cetyltrimethylammonium bromide (CTAB) protocol described by Cullings (1992) and Xiang et al. (1998). DNA quantity was analyzed with Qubit 2.0 (Life Technologies, USA) and quality was measured using a NanoDrop spectrophotometer 2000 (ThermoFisher Scientific, USA) and 1% w/v agarose gels. Sequencing was performed at the High-Throughput Sequencing Facility (HTSF) at UNC Chapel Hill. For Illumina sequence libraries (Illumina, CA, USA), ~250 ng of total DNA was utilized. Agilent 2100 Bioanalyzer (Agilent Technologies, USA) was used to select ~450 bp fragments for Novaseq 6000, 250 bp paired-end (PE) sequencing. For the Oxford Nanopore sequencing, ~2000 ng of high-molecular weight DNA was prepared using the ligation sequencing kit (SQK-LSK109) and sequenced on two R9.4.1 flowcells (Oxford Nanopore Technologies, Oxford, UK).

### 2.2 ptGAUL pipeline and validation

We generated a pipeline to facilitate plastome assembly using long-read data, which can be applied to both PacBio and ONT raw reads. The ptGAUL pipeline (Figure 1) includes three major parts: filtering long reads, setting the depth of coverage, and assembling the filtered plastid data. Step 1: use minimap2 (Li et al., 2018) to find all reads that map partially or completely to the closely related reference plastome, followed by filtering all reads using a customized bash script. Then, use seqkit (Shen et al., 2016) to keep long reads greater than a specified length (default is 3000 bp, “-f” in ptGAUL). Step 2: calculate the coverage by assembly-stats (available in https://github.com/sanger-pathogens/assembly-stats). If the coverage is over 50×, apply seqtk (available in https://github.com/lh3/seqtk) to randomly select a subset of data including about 50× coverage of the plastome (higher coverage might fail in assembly). Step 3: use Flye (Kolmogorov et al., 2019) to assemble the plastome. When only three contigs were detected in the graphical fragment assembly (gfa) file, we used combined_gfa.py, a customized python script, to assemble the plastome into two different paths. Otherwise, the assembly result was checked manually using Bandage. All pipeline code was deposited on Github (https://github.com/Bean061/ptgaul). Step 3: if ptGAUL was different from the assembly coverage setting in Flye (--asm-coverage), ptGAUL implements seqtk to randomly choose a subset of long reads to minimize the bias of read selection. After assembly, if short-read sequencing data are available, the FM-index Long Read Corrector (FMLRC) software (Wang et al., 2018) is recommended to polish and improve the accuracy of the final assembled sequences because it can generate more accurate assembly result (Mak et al., 2022). All analyses related to ptGAUL were run using 10 CPUs and 40G RAM on the longleaf cluster at UNC Chapel Hill.

**Figure 1.**
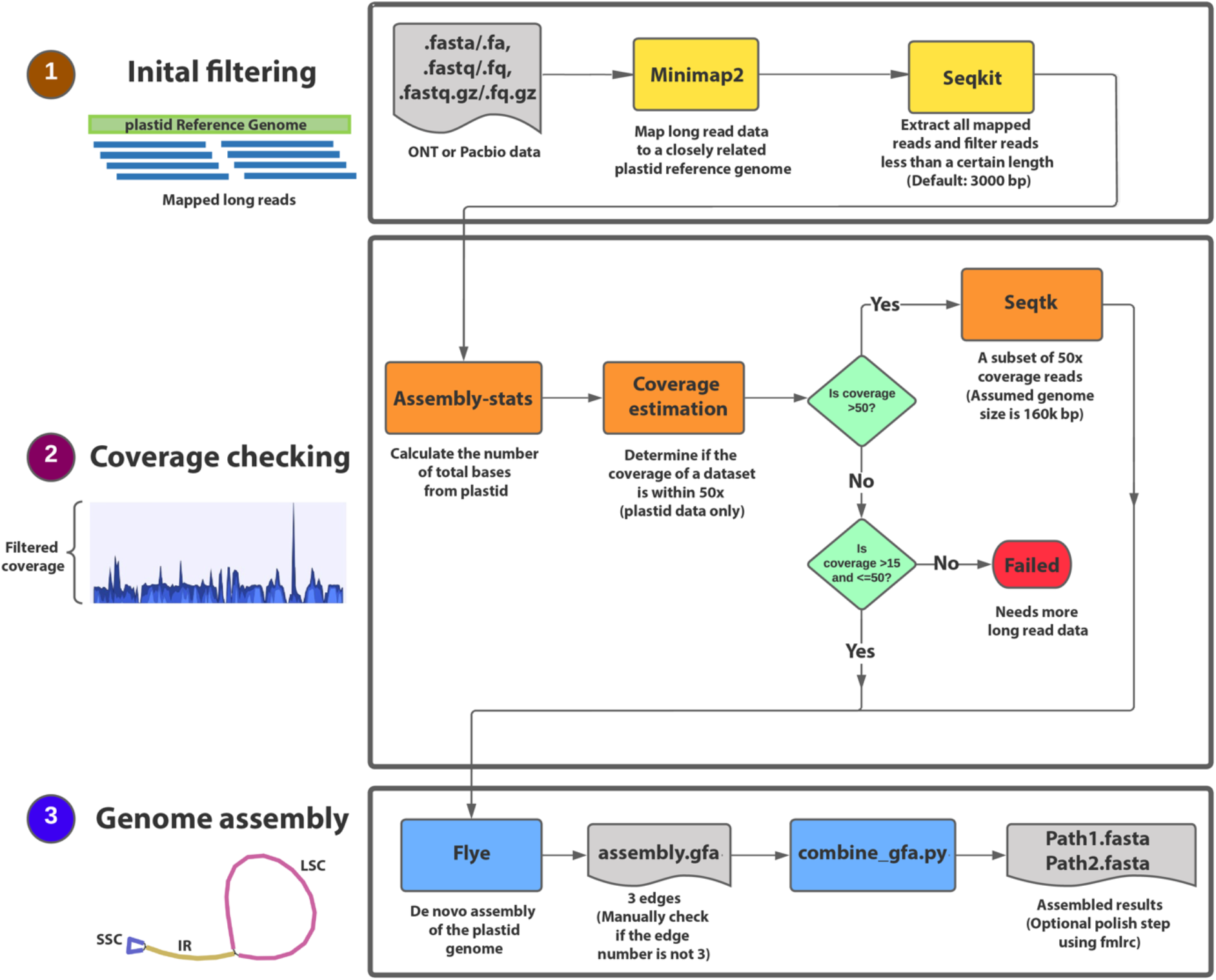
ptGAUL workflow. The program starts with an initial filtering step to filter the long reads of the target species using at least one closely related reference plastome (1). Subsequently, the coverage for those filtered long reads is calculated and filtered to make sure it is about 50× (2). Finally, two paths of plastomes were obtained through Flye and a customized python script, combine_gfa.py (3).

Long-read data from 16 published plastomes in NCBI were used to validate the efficacy of ptGAUL for assembly (Table 1). Comparative analyses were conducted including the number of assembled contigs, total genome size (bp) and nucleotide sequence identity between the published results and those obtained with ptGAUL (pairwise identity in alignment) using Geneious v.2022.2 (Kearse et al., 2012).

**Table 1.** ptGAUL performance on 15 published sequence data sets, including the information of assembled plastome from published papers and the information on assembled plastomes from ptGAUL. NA means low nucleotide sequence identity between assembled plastome between published data and our data. S means the samples are well assembled by ptGAUL, while F means the samples failed using ptGAUL. * column includes the references we used for genome assembly in ptGAUL and the bold references were considered as references for comparisons with ptGAUL results.

| Species | Library preparation and sequencing methods | Raw read No./N50 (bp) | Reference | Plastid size from ptGAUL (bp) (% nucleotide sequence identity to references) | Number of assembled plastid contigs from ptGAUL | Plastome reference used for ptGAUL (reference length from original studies)* |
| --- | --- | --- | --- | --- | --- | --- |
| *Arctostaphylos glauca* | WGS/PacBio | 1814591/15245 | Huang et al., 2022 | 150241 (NA) | 3 (S) | NC_035584.1/NC_042713.1/NC_047438.1/**JAHSPW020000272.1 (118663 bp)** |
| *Lepidium sativum* | WGS/PacBio | 400322/7277 | Zhu et al., 2019 | 153666 (99.9%) | 3 (S) | **NC_047178.1 (154997 bp)** |
| *Chaetoceros muellerii* | WGS/PacBio | 87313/12921 | Li & Deng, 2021 | 117304 (99.8%) | 1 (S) | **MW004650.1 (116284 bp)** |
| *Potentilla micrantha* | WGS/PacBio | 28638/2464 | Ferrarini et al., 2013 | 159850 (99.8%) | 3 (S) | **NC_015206.1 (155691 bp)** |
| *Durio zibethinus* | WGS/PacBio | 853182/9670 | Shearman et al., 2020 | 142806 (99.95%) | 1 (S) | **MT321069 (163974 bp)** |
| *Beta vulgaris* | WGS/PacBio | 96874/3980 | Stadermann et al., 2015 | 155383 (99.9%) | 3 (S) | **KR230391.1 (149722 bp)** |
| *Eleocharis dulcis* | WGS/PacBio | 68167/16288 | Lee et al., 2020 | 199919 (99.5%) | 3 (S) | **NC_047447.1 (199561 bp)** |
| *Eucalyptus pauciflora* | WGS/ONT | 705554/24988 | Wang et al., 2018 | 158561 (99.0%) | 1 (S) | MZ670598.1/HM347959.1/NC_014570.1/AY780259.1/**NC_039597.1 (159942 bp)** |
| *Leucanthemum vulgare* | Long range PCR/ONT | 18031/7900 | Scheunert et al., 2020 | 119593 (NA) | 5 (F) | **NC_047460.1 (150191 bp)** |
| *Oryza glaberrima* | Plastid capture/ONT | 81363/4319 | Bethune et al., 2019 | 124133 (NA) | 4 (F) | **NC_024175.1 (132629 bp)** |
| *Cenchrus americanus* | Plastid capture /ONT | 105760/5580 | Bethune et al., 2019 | 143162 (96.6%) | 3 (S) | **NC_024171.1 (140718 bp)** |
| *Digitaria exilis* | Plastid capture /ONT | 141250/4028 | Bethune et al., 2019 | 136650 (96.0%) | 3 (S) | **NC_024176.1 (140908 bp)** |
| *Podococcus acaulis* | Plastid capture /ONT | 249417/2621 | Bethune et al., 2019 | 81976 (NA) | 2 (F) | **NC_027276.1 (157688 bp)** |
| *Raphia textilis* | Plastid capture /ONT | 83833/2495 | Bethune et al., 2019 | 60089 (NA) | 2 (F) | **NC_020365.1 (157270 bp)** |
| *Phytalephas aequatorialis* | Plastid capture /ONT | 202925/2551 | Bethune et al., 2019 | NA | (F) | **NC_029957.1 (159075 bp)** |
| *Picea glauca* | WGS/PacBio | 563675/4671 | Soorni et al., 2017 | 123476 (98.9%) | 1 (S) | **NC_021456.1 (124084 bp)** |

### 2.3 Assembly and comparison of four *Juncus* species

The Illumina Novaseq 6000 platform (Illumina, USA) was used to generate 250 bp, paired-end (PE) reads for *Juncus roemerianus* and *J. validus*. Reads were *de novo* assembled using GetOrganelle v1.7.5 (Jin et al., 2020) with default settings. Long-read data were also generated using ONT for *J. roemerianus* and *J. validus*. Long-read data were assembled using ptGAUL with default (3000 bp) filtering parameters (“-f”) and using all eight *Juncus* plastomes on GenBank (Table 2) as references for the filtering step. We verified the assembly graph results (gfa file) from Flye using the visualization in Bandage v 0.8.1 (Wick et al., 2015). Then, we conducted FMLRC to polish the final plastomes (an optional step in the ptGAUL pipeline). To examine the assembly result, we mapped all raw Illumina reads and raw ONT reads of each *Juncus* species to our polished assembly and tested the evenness of the coverage at all sites. If every site shares a similar coverage of raw reads without gaps in coverage, this usually indicates a good *de novo* assembly result. We used the samtools v.1.9 (Danecek et al., 2021) depth function to record read depth at every site, followed by a dot plot created by the matplotlib library (Hunter, 2007) in python. We downloaded the raw whole genome sequencing data of *J. effusus* and *J. inflexus* (both ONT reads: SRR14298760 and SRR14298751 and Illumina reads: SRR14298746 and SRR14298745 from Planta et al., 2022) to assemble the plastomes following the same steps above.

**Table 2.** Summary of features of the plastid genomes of four *Juncus* species, including length, GC content, and gene numbers. The plastome data of *Juncus effusus* were from two different two different sources, this paper and Lu et al. (2021). PCG =protein-coding genes.

| Genome Features | *J. effusus* | *J. effusus* | *J. inflexus* | *J. roemerianus* | *J. validus* |
| --- | --- | --- | --- | --- | --- |
| Accession No. | NC_059754.1 | Present study | Present study | OP235509 | OP235510 |
| No. of Illumina read clusters | 12443053 | 96,653,565 | 83412073 | 158922322 | 156712430 |
| No. of ONT reads and N50 | 0 | 2960380/21529 | 2735792/24397 | 427549/15998 | 243884/14365 |
| Genome size (bp) | 170612 | 178158 | 181566 | 196852 | 147183 |
| LSC length (bp) | 81818 | 86497 | 86649 | 82944 | 87215 |
| SSC length (bp) | 7522 | 7539 | 7509 | 7902 | 2046 |
| IR length (bp) | 40636 | 42061 | 43704 | 53003 | 28961 |
| Overall GC content, % | 36.0 | 35.9 | 35.6 | 32.2 | 34.7 |
| GC content in LSC, % | 33.2 | 33.2 | 33.3 | 33.1 | 31.6 |
| GC content in SSC, % | 26.3 | 26 | 26.2 | 26.5 | 23 |
| GC content in IR, % | 39.7 | 39.5 | 38.7 | 37.5 | 39.8 |
| Total No. of genes | 129 | 133 | 134 | 136 | 114 |
| No. of unique genes | 105 | 106 | 106 | 106 | 93 |
| No. of unique PCGs | 72 | 72 | 72 | 72 | 60 |
| No. of unique tRNA genes | 29 | 30 | 30 | 30 | 29 |
| No. of unique rRNA genes | 4 | 4 | 4 | 4 | 4 |

After assembly, we uploaded plastomes of four *Juncus* species (*J. roemerianus, J. validus, J. effusus*, and *J. inflexus*) to GeSeq online (Tillich et al., 2017) for annotation using Chole (Zhong, 2020), HMMER (Finn et al., 2011) and BLAT (Kent, 2002). We manually checked the start and stop codons of each annotated gene using Geneious v.2022.2. The genes not in frame in each *Juncus* species were either adjusted or removed after a careful comparison with *Typha latifolia* plastid annotation (NC_013823; Guisinger et al., 2010) by mapping the annotations to our *Juncus* assemblies. For the uncertain tRNAs, we confirmed the tRNA secondary structures via RNAfold WebServer (Hofacker, 2003). Linear plastome maps were drawn with OGDRAW v. 1.2 (Lohse et al., 2013). Circular representations were drawn using Circoletto (Darzentas, 2010) to visualize the repeats.

### 2.4 Examination of repeats and rearrangement events in *Juncus*

We removed one copy of the IR region prior to repeat analyses to avoid counting the repeats from IR region. We implemented BLAST v.2.8.1+ (Altschul et al., 1990) and Tandem Repeats finder v4.09.1 (Benson, 1999) to detect the dispersed repeats and tandem repeats, respectively, following the steps from Lee et al. (2020). We manually checked the result and eliminated duplicated blast hits and recorded the total number of distinct dispersed repeats. We also downloaded complete plastomes of *Eriocaulon decemflorum* (NC_044895; Darshetkar et al., 2019) and two early diverging Poales, *Typha latifolia* (NC_013823; Guisinger et al., 2010) and *Ananas comosus* (NC_026220; Nashima et al., 2015), for comparison. All the plots were drawn using matplotlib (Hunter, 2007) in python.

We focused on the four confirmed assemblies of *Juncus*, i.e., *J. roemerianus, J. validus, J. effusus*, and *J. inflexus* for characterizing and comparing the rearrangements in *Juncus* plastomes. The other eight publicly available (Lu et al., 2021; Wu et al., 2021) *Juncus* plastomes on GenBank were excluded from the rearrangement analyses (Table 2) because of the uncertain assemblies resulting from short-read data. To eliminate uncertainty in short-read assemblies, we compared them to the *J. effusus* plastome assembled from short-read data (Lu et al., 2021) and to the ptGAUL-assembled plastome of *J. effusus* from long-read data (Planta et al. 2022). To detect rearrangement events within *Juncus*, whole-genome alignments of *J. roemerianus*, *J. validus*, *J. effusus*, and *J. inflexus* were performed to examine the arrangement of locally colinear blocks (LCBs) using progressiveMauve (Darling et al., 2004). One copy of the IR was removed from plastomes prior to Mauve alignment to prevent spurious alignments. *Typha latifolia* was employed as a reference and *Ananas comosus* and *Eriocaulon decemflorum* were also included.

## 3 RESULTS

### 3.1 Validation of ptGAUL

Overall, ptGAUL assemblies were successful; assemblies contained either one or three contigs in 11 of 16 the species, with plastome sizes similar to those reported previously (indicated with “S” in Table 1). The assembly graph results (gfa files) indicating plastome structure were visualized and confirmed by Bandage and deposited in Github (https://github.com/Bean061/ptgaul). Assembled plastomes had > 95% nucleotide sequence identity to the references, however the plastome of *Arctostaphylos glauca* was 31,578 bp longer (21% total length) than the published data (Table 1). ptGAUL failed to assemble plastomes of five species (indicated with F in Table 1). The ptGAUL pipeline produced consistent and reliable results when provided with a dataset of long reads (> 5000 bp N50) with ~50X coverage of the plastome.

Our results indicated that different library preparations affected plastome assembly, regardless of the long-read sequencing platform (PacBio or ONT) employed (Table 1). Plastomes derived from a whole genomic sequencing approach assembled correctly (either one or three contigs), with a reasonable plastome length and structure (by Bandage), while the plastomes using plastid capture approaches (i.e., long range PCR and long fragment target capture) were more fragmented and had a smaller genome size. For example, *Leucanthemum vulgare* had a similar N50 value to *Lepidium sativum* (7900 bp and 7277 bp, respectively), but the *Leucanthemum vulgare* library prepared using long range PCR failed in plastome assembly. All five failed datasets involved the plastid capture approach and most of the raw sequence reads had relatively short length with small N50 values (less than 5000 bp) (Table 1).

### 3.2 Plastome features of four *Juncus* species

We generated 158,922,322 and 156,712,430 short reads for *Juncus roemerianus* and *J. validus*, respectively along with 427,549 ONT reads from *J. roemerianus* (N50 value: 15,998 bp) and 243,884 ONT reads from *J. validus* (N50 value: 14,365) (Table 2). The data were deposited at NCBI with BioProject accession: PRJNA865266. We also downloaded sequence data (PRJNA723756)of *J. effusus* and *J. inflexus* from Planta et al. (2022) (Table 2). The ptGAUL pipeline produced three contigs each for *J. validus* and *J. roemerianus* (Figure S1a, b) sequenced in this study, and one contig each for *J. effusus* and *J. inflexus* sequenced by Planta et al. (2022) (Figure S1c, d). The final assembled plastomes of *J. validus, J. roemerianus, J. effusus*, and *J. inflexus* ranged from 147,183 to 196,852 bp, had similar sized LSCs from, different sizes of the SSC from, and large differences in IR size (Table 2).

The assemblies for the four *Juncus* species were verified by mapping both Illumina and ONT reads back to the assembly. All mapping results showed a high and even coverage of both species (Figure 2 c-f; Figure S2 a,b,d,e). There were no gaps in assemblies regardless of sequencing platform. Annotation of the four *Juncus* plastomes revealed that they contained 114 to 136 genes, 93 to 106 of which were unique. There were 60 to 72 unique protein coding genes (PCGs), 29-30 tRNA genes, and four rRNA genes (Table 2). *Juncus roemerianus* has the greatest gene number at 136, which is similar to the *J. effusus* (133), *J. inflexus* (134), and basal Poales ancestors *Typha latifolia* (133), *Ananas comosus* (132), *Eriocaulon decemflorum* (135) (Table S2). *Juncus effusus* and *J. inflexus* shared a highly similar gene content while *J. validus* lacks 11 *ndh* genes, *rps15* and *trnT-GGU* (Table S2). (Figure S3; Table S2).

**Figure 2.**
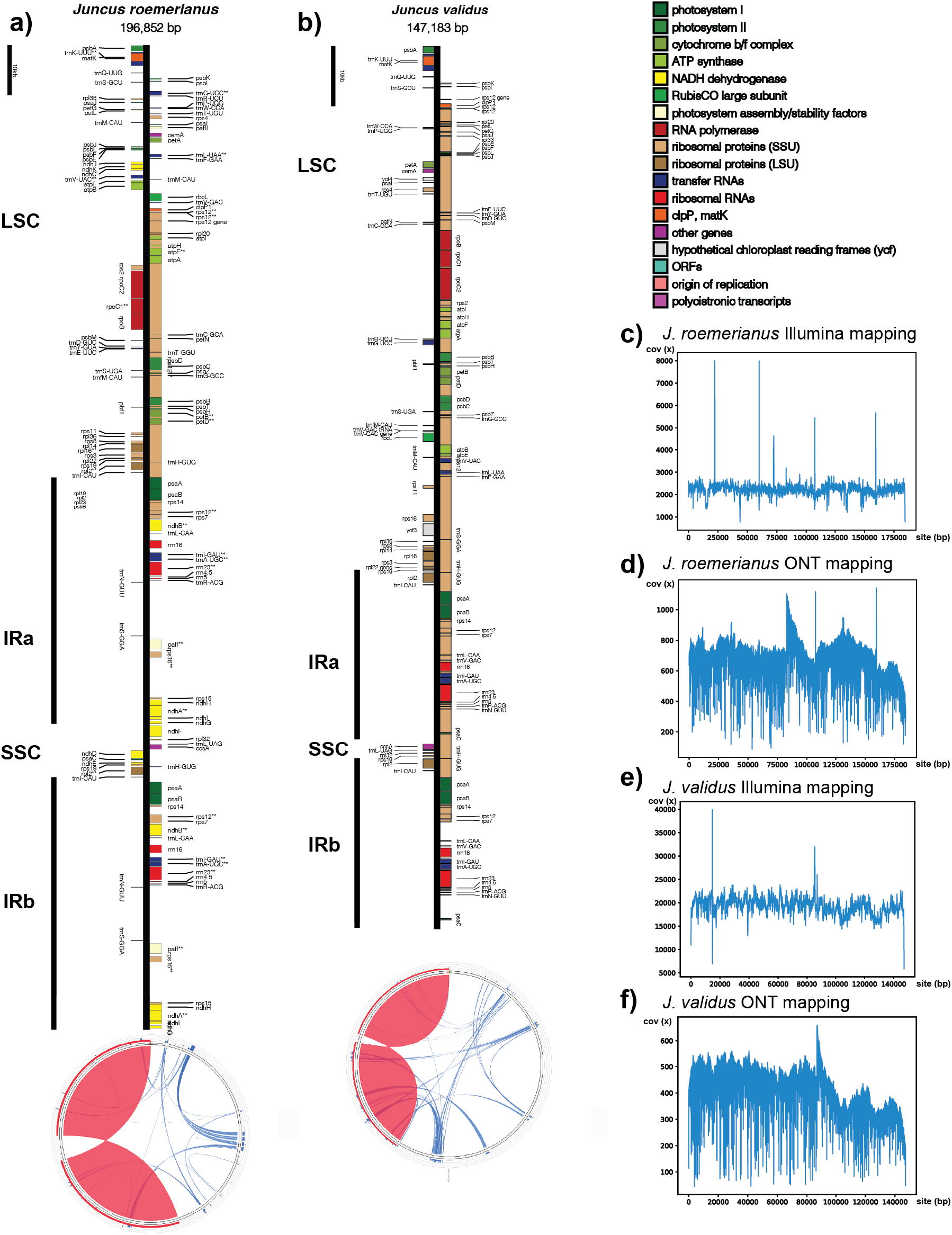
Plastome structural maps and read coverage graphs of *J. roemerianus* and *J. validus*. a) and b) Linear maps of *J. roemerianus* and *J. validus* plastome, respectively, were drawn by OGDRAW (Lohse et al., 2013). Genes that belong to different functional groups are color-coded. Small single copy (SSC), large single copy (LSC), and inverted repeats (IRa, IRb) are indicated for both plastomes. Circular representatioins of the two *Juncus* plasstomes were used to shot locations of repetitive DNA using Circoletto (Darzentas, 2010). The blue lines represent dispersed repeats in the plastome, while red regions represent the IR regions. c) - f) Read coverage plots of *J. roemerianus* and *J. validus* using Illumina reads and ONT reads,respectively, showing the good quality of the assemblies. The *x* axis represents the position in the plastome, while *y* axis represents the coverage.

### 3.3 Verification of published *Juncus effusus* plastome

We compared the *J. effusus* published assembly based on short-read data (Lu et al., 2021, MW366789) with our new assembly using ptGAUL of long-read data of Planta et al. (2022). The result indicated that the short-read assembly generated by Lu et al. was >7.5 kb shorter than our long-read assembly (170,612 bp versus 178,158 bp). The mapping results showed that our assembly was well supported by both long-read and short-read data from Planta et al. (2022) (Figure S2a,b), yet unsupported by the Illumina reads from Lu et al. (2021) with 777 positions with less than 10x coverage, including 295 positions that have no read coverage (Figure S2c). The previous short-read assembly of *J. effusus* (MW366789) was not supported by the long-read data from Planta et al. (2022). Based on this result, we removed the eight publicly available *Juncus* plastomes assembled with short-read data prior to the comparative analyses of plastomes.

### 3.4 Repeats in *Juncus* plastomes

Repeat analyses identified many dispersed and tandem repeats in the four *Juncus* plastomes (17.2 – 24.3 % of genome without IRa) in comparison with basal Poales and *Eriocaulon* (1.8 – 3.3 % of genome without IRa) (Table 3). The combined length of both dispersed and tandem repeats in *Juncus* plastomes ranged from 22,577 bp (*J*. *validus)* to 34,027 bp (*J*. *roemerianus*), which was far greater than *Typha* (4,436 bp), *Ananas* (3,552 bp) and *Eriocaulon* (2,227 bp) (Table 3). When dispersed repeats were parsed into five different size classes, *Juncus* plastomes contained a greatly increased number of dispersed repeats than basal Poales and *Eriocaulon* (Figure 3 and Table S3). Larger repeats (>201 bp) were found only in *Juncus* (Figure 3 and Table S3). Among four *Juncus* plastomes, *J*. *effusus* and *J*. *validus* had more abundant dispersed repeats yet *J*. *roemerianus* was the only one with a repeat larger than 1 kb. *Juncus* plastomes also experienced substantial accumulation of tandem repeats (Table 3). Tandem repeat accumulation was higher than that of dispersed repeats in *J*. *inflexus* and *J*. *roemerianus*. All four *Juncus* plastomes contained exceptionally expanded tandem repeats, ranging from 4.6 – 6.6 kb, some of which contain *clpP* (Table S4).

**Figure 3.**
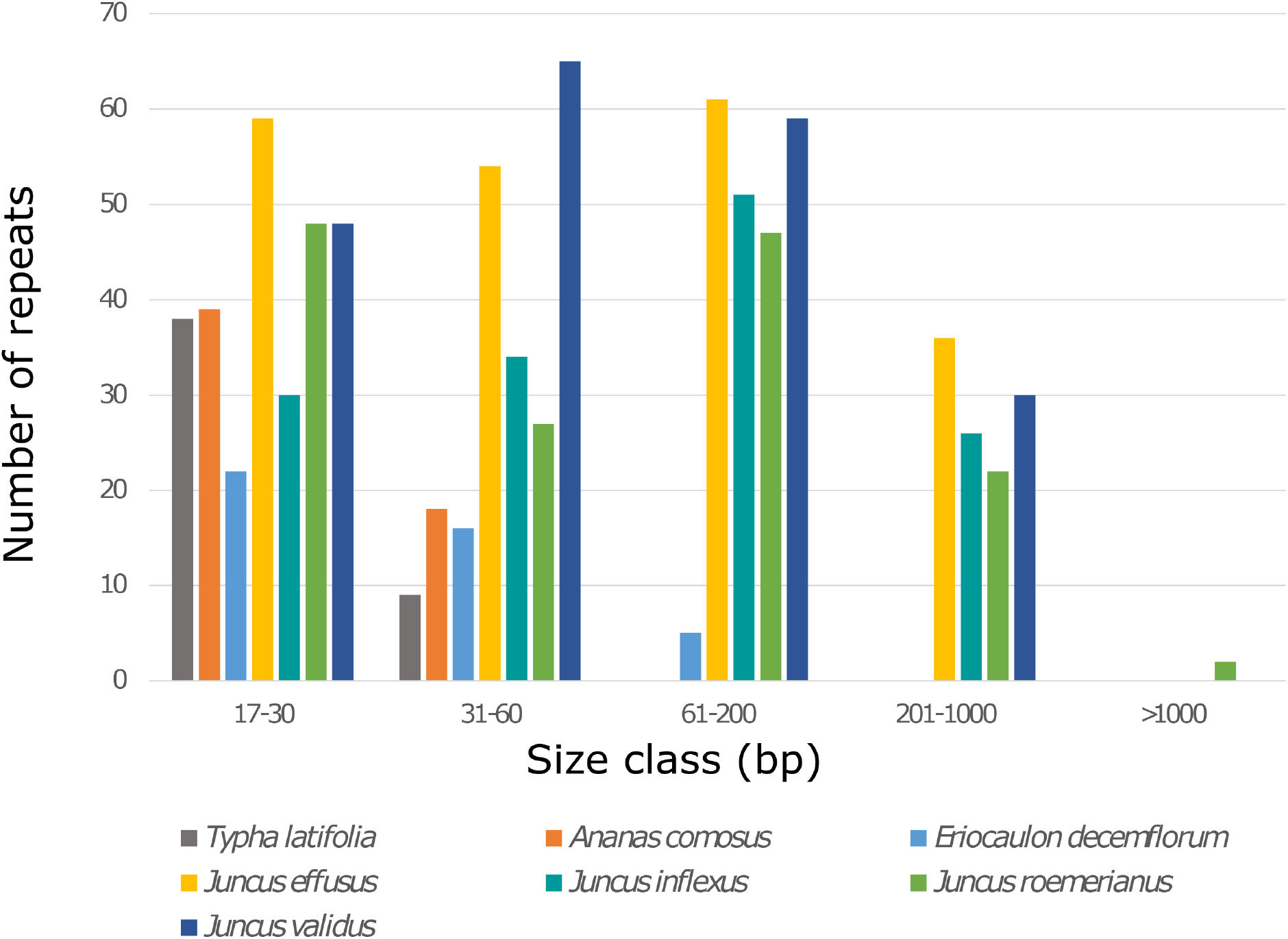
Bar plot of dispersed repeats in plastomes from seven Poales species, including four newly assembled *Juncus* species.

**Table 3.** Statistics of dispersed and tandem repeats in *Typha, Ananas, Eriocaulon*, and *Juncus* plastomes

| Species | *Typha*<br>*latifolia* | *Ananas*<br>*comosus* | *Eriocaulon*<br>*decemflorum* | *Juncus*<br>*effusus* | *Juncus*<br>*inflexus* | *Juncus*<br>*roemerianus* | *Juncus validus* |
| --- | --- | --- | --- | --- | --- | --- | --- |
| Genome size (no IRa) | 134,642 | 132,862 | 125,164 | 136,097 | 137,859 | 143,849 | 118,221 |
| GC % | 35.5 | 36.3 | 34.2 | 34.8 | 34.6 | 34.3 | 33.5 |
| **Dispersed repeat (DR)** |  |  |  |  |  |  |  |
| Length of DRs | 1,210 | 1,495 | 1,418 | 15,117 | 13,229 | 14,712 | 14,714 |
| GC % | 33.7 | 36.3 | 33 | 35 | 36 | 35.7 | 34 |
| GC % without DR | 35.5 | 36.3 | 34.2 | 34.7 | 34.6 | 34.1 | 33.3 |
| % of DR in genome | 0.9 | 1.1 | 1.1 | 11.1 | 9.6 | 10.2 | 12.4 |
| **Tandem repeat (TR)** |  |  |  |  |  |  |  |
| Length of TRs | 3,270 | 2,057 | 859 | 12,248 | 15,783 | 22,978 | 8,797 |
| GC % of TRs | 8.8 | 18.4 | 20 | 34.2 | 33.4 | 32.1 | 32.1 |
| Genome size without TRs | 131,372 | 130,805 | 124,305 | 123,849 | 122,076 | 120,871 | 109,424 |
| GC % without TRs | 36 | 36.6 | 34.3 | 34.8 | 34.8 | 34.8 | 33.6 |
| % of TRs in genome | 2.4 | 1.5 | 0.7 | 9.0 | 11.4 | 16.0 | 7.4 |
| **Total repeat** |  |  |  |  |  |  |  |
| Length of total repeats | 4,436 | 3,552 | 2,227 | 23,345 | 26,451 | 35,027 | 22,577 |
| GC % of total repeats | 12.6 | 21.5 | 27.5 | 34.7 | 34.7 | 33.6 | 33.5 |
| GC % without total repeats | 36 | 36.6 | 34.3 | 34.8 | 34.8 | 34.5 | 33.4 |
| % of total repeats in genome | 3.3 | 2.7 | 1.8 | 17.2 | 19.2 | 24.3 | 19.1 |

### 3.5 Rearrangement of *Juncus* plastomes

Whole-genome alignment using progressiveMauve (Figure 4) detected 27 LCBs from seven complete plastomes (*Typha latifolia*, *Ananas comosus*, *Eriocaulon decemflorum*, *Juncus effusus*, *J. inflexus*, *J. roemerianus*, and *J. validus*). The plastomes of the two basal Poales and *Eriocaulon* were colinear, whereas all *Juncus* species have many breakpoints (BP) relative to the reference, *T. latifolia* (Figure 4; Table 4). When compared with basal Poales plastomes, the BP and reversal distances were 15 and 19, in *J. effusus* and *J. inflexus*, repspectively. *Juncus roemerianus* has the largest BP (17) and reversal distances (20), and *J. validus* has the smallest BP (14) and reversal distances (17). Among the four *Juncus*, 27 LCBs were identified (Figure S4). While *J. effusus* and *J. inflexus* shared the same gene order, widespread rearrangements were detected in the other two species (*J. roemerianus* and *J. validus*).

**Figure 4.**
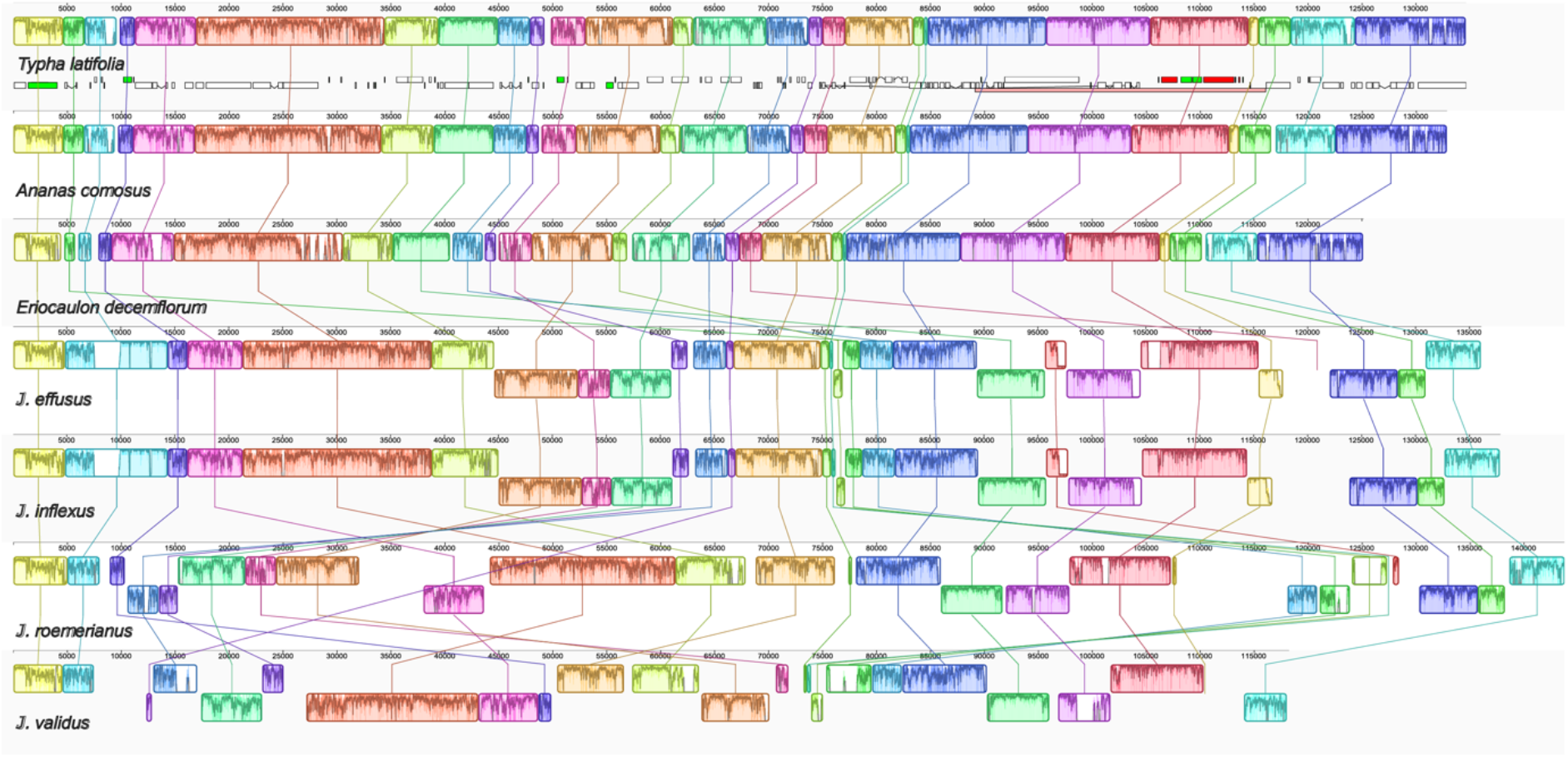
Whole plastome alignment of seven Poales species, including four newly assembly *Juncus* and *Typha latifolia, Ananas comosus*, and *Eriocaulon decemflorum*. The local colinear blocks (LCBs) were identified by progressiveMauve with *Typha* plastome as the reference. The corresponding LCBs among seven plastomes are shaded and connected with a line of the same color. LCBs that are flipped indicate inversions. Numbers on the upper *x*-axis are genome map coordinates in basepairs (bp).

**Table 4.**
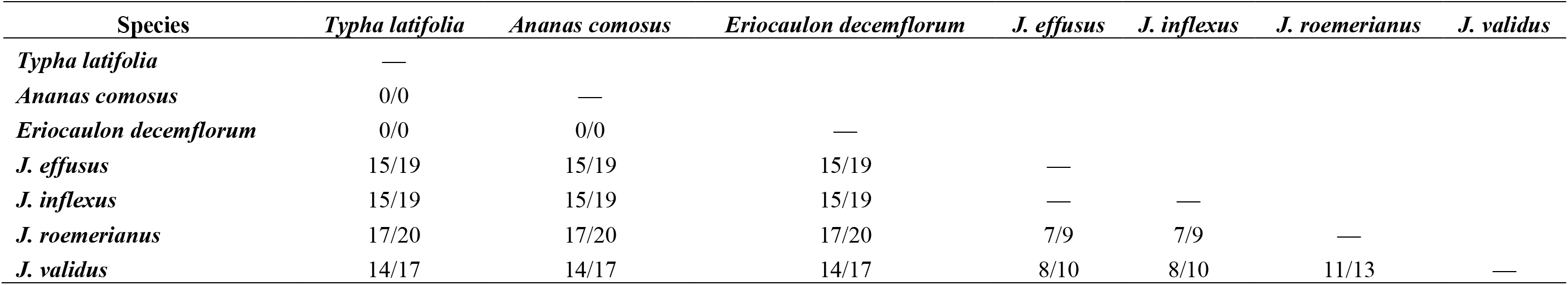
Summary of breakpoint and reversal distances for plastomes of *Juncus, Eriocaulon* and basal Poales.

| Species | *Typha latifolia* | *Ananas comosus* | *Eriocaulon decemflorum* | *J. effusus* | *J. inflexus* | *J. roemerianus* | *J. validus* |
| --- | --- | --- | --- | --- | --- | --- | --- |
| *Typha latifolia* | — |  |  |  |  |  |  |
| *Ananas comosus* | 0/0 | — |  |  |  |  |  |
| *Eriocaulon decemflorum* | 0/0 | 0/0 | — |  |  |  |  |
| *J. effusus* | 15/19 | 15/19 | 15/19 | — |  |  |  |
| *J. inflexus* | 15/19 | 15/19 | 15/19 | — | — |  |  |
| *J. roemerianus* | 17/20 | 17/20 | 17/20 | 7/9 | 7/9 | — |  |
| *J. validus* | 14/17 | 14/17 | 14/17 | 8/10 | 8/10 | 11/13 | — |

## 4 DISCUSSION

### 4.1 ptGAUL application and suggestions for sequencing approach

The ptGAUL pipeline generated either one or three contig(s) for 11 publicly available datasets using either PacBio or Oxford Nanopore data (Table 1). However, it failed to assemble the data from five species generating more than three short contigs and predicted much smaller plastome size, which is less than optimal (Table 1). In successful cases, the assemblies were highly similar to the published short-read assemblies with over 96-99% nucleotide sequence identity. The lower percent identity between *Cenchrus americanus* and *Digitaria exilis* and their reference assemblies may be due to different sequencing approaches between the Mariac et al. (2014) combined plastid capture method and Illumina sequencing and our long-read approach. For *Arctostaphylos glauca*, we used the read mapping method to verify that our assembly was more reliable than the result of Huang et al. (2022) as it showed more proportional coverage across the entire plastome (Figure S5). This difference could be caused by the selection of a distantly-related reference genome (*Camellia taliensis*) from another family by Huang et al. (2022).

We found that the five failed samples had some features in common. For example, the sequencing approaches in failed assemblies were different from the whole genomic sequencing method of those that were successful. In the *Leucanthemum vulgare* study, long-range PCR was implemented to generate amplicons that were then sequenced to produce a set of long reads that had an N50 value of ~ 8000 bp (Scheunert et al., 2020). In the remaining failed assemblies, plastid capture was utilized (Bethune et al., 2019). The PCR processes in both studies can greatly increase the bias among different plastome regions, e.g. AT- and GC-rich regions do not amplify as efficiently as other regions (Quail et al., 2012). This could lead to underrepresentation/unevenness in read coverage of different regions resulting in many fragmented assemblies/contigs. Furthermore, the probes were designed based on the plastome data from distantly related species (Bethune et al., 2019), which may be unable to capture all plastome fragments for the target non-model species due to the divergence between the probe regions and the genome being captured. Additionally, the sequences obtained from PCR methods tend to be much shorter than the reads generated from sequencing total genomic DNA (see N50 values in Table 1). The low N50 values could also result from degraded DNA from poor storage, use of silica dried or herbarium material and/or DNA extraction method. For example, the Qiagen DNEasy Plant kits can generate high quality DNA for short-read sequences because the column shreds the DNA to a maximum of ~25 Kb fragments (Qiagen, 2006). CTAB, SDS or other methods that can produce much higher molecular weight (HMW) DNA are preferred for third generation sequencing (Mayjonade et al., 2016; Jung et al., 2019), emphasizing the importance of sample preparation. Likewise, the assembly approaches, parameter combinations, read coverage, and the presence of nuclear genome and/or mitogenome contaminants could impact the completeness of an assembly (Jung et al., 2019; Scheunert et al., 2020).

Overall, considering the read length and read coverage, ptGAUL performs well for HMW samples using total genomic sequencing resulting in high N50 values. Therefore, we recommend using HMW DNA extraction methods to isolate highly intact DNA, followed by long-read sequencing and subsequent assembly using ptGAUL.

### 4.2 Long-read data for plastome assembly

We found that short-read data alone may be insufficient to accurately assemble plastomes in species with many long dispersed repeats. This phenomenon has been seen in several lineages including *Eleocharis* (Lee et al., 2020) and *Monsonia* (Ruhlman et al., 2017). Plastome assembly using GetOrganelle for 11 *Juncus* species (12 accessions) failed using Illumina short reads only, including two samples in this study (Fig. S6). All *Juncus* plastome assemblies indicated either many fragmented contigs or many assembly paths (Figure S6). This is because the many long dispersed repeats in *Juncus* plastomes are longer than the kmer size/length of short reads. Based on our *J. effusus* plastome comparison, the final assembly length and total number of genes based on short read data is much shorter than the ones assembled from long read data (Table 2, Table S2), which might be caused by the random selection one of the paths as the final assembly when using short read data. Other studies demonstrated that this issue can be resolved by a three-step approach: comparing different contigs from short-read assemblers (e.g. SPADES, Velvet), manually checking through the contigs compared with closely related species, and long range PCR to confirm assemblies (Lee et al., 2020; Ruhlman et al., 2017). This approach requires considerable time and effort.

Our ONT data resolved the plastome structure of four *Juncus*, confirming prior work (Lee et al., 2020; Ruhlman et al., 2017) showing that long-read data vastly improves assembly of the plastomes with many long repeats. Based on our study and that of Scheunert et al. (2020) ~50X mapping coverage of long-read data can result in an accurate plastome assembly. In our study, long reads of plastid origin represented 5%-6% of reads generated from total genomic DNA of *Juncus*. Assuming 5% plastid DNA content from whole-genome HMW extractions, to generate ~ 50X coverage of a 160,000 bp plastome requires only ~160 Mbp reads per sample. Currently one chip of ONT generates ~10 Gbp of sequence data, enabling multiplexing up to 64 samples at a consumables cost of ~$1000 USD (based on the price from HTSF at UNC Chapel Hill).

Although several assembly tools have been developed, several issues persist. Some pipelines/software are no longer maintained (i.e., Sprai, Celera Assembler, Organelle_PBA). The assemblers of Syme et al. (2021); others, Canu, and Hinge (Wang et al., 2018) cannot generate a consistent plastome assembly result with one contig when using different coverages of data. Unicycler (Wick et al., 2017) is computationally intensive and does not produce well resolved assemblies when dealing with complicated plastomes with many long repeats. Compared to current published pipelines for plastome assembly, ptGAUL can help generate an accurate plastome assemblies in less than ~20 minutes (10 CPUs and 40G RAM), making it highly convenient. Thus, ptGAUL should greatly facilitate plastome assembly of long-read data for phylogenetic and molecular evolutionary studies, especially in plastomes with a significant fraction of long repeat regions. Although ptGAUL can expedite plastome assembly, researchers still need to pay close attention to the species with multiple plastid types, such as *Eleocharis* (Lee et al., 2020) and *Monsonia* (Ruhlman et al., 2017).

### 4.3 *Juncus* plastome organization

While many Poales genera contain plastomes with conserved gene order and content (Jones et al., 2007), including *Typha* (Guisinger et al., 2010), *Ananas* (Redwan et al., 2015) and *Eriocaulon* (Darshetkar et al., 2019), the data from the four *Juncus* examined here suggest that at least some species in this group contain plastome features atypical to most angiosperms. A limited number of complete plastome sequences are available from *Juncus* or other Juncaceae, but recently assemblies of two *Eleocharis* plastomes, in the sister family Cyperaceae (Hochbach et al., 2018), revealed accumulated duplications, gene losses, gene order rearrangements and intraindividual structural heteroplasmy (Lee et al., 2020). Similar phenomena contributed to size variation in the four *Juncus* plastomes, which ranged from 147,183 bp to 196,852 bp (Table 2). Many long repeats, including an unusually high number of dispersed repeats of 61 – 200 bp and 201 – 1000 bp, were present in the four *Juncus* with the greatest accumulation in *J. effusus*. Repeats >1000 bp were detected only in *J. roemerianus* (Table S3; Figure 3). Accumulation of large repeats may predispose plastome rearrangements in addition to contributing to overall size expansion (Tables 2–3, Figure 4) yet at present it is not clear if repeat accumulation predicated rearrangement or *vice versa* (Lee et al., 2021).

Similar repeat accumulation and plastome rearrangement occur in other taxonomic groups. In the *Trachelium caeruleum*, there are gene-order changes, along with gene duplication, pseudogenization and loss were identified, as well as an abundance of variously sized repeats (Haberle et al., 2008). A relationship between repeat accumulation and rearrangement was suggested (Kim & Lee, 2005); studies of *Pelargonium* (Chumley et al., 2006), *Jasminum, Mendora* (Lee et al., 2007) and *Trifolum* (Cai et al., 2008) plastomes show early support for the theory. Many of the repeated sequences, when plotted onto the assembled plastid chromosomes, clustered at rearrangement endpoints. The relationship is also supported by findings in bacterial genomes where repeated sequences lead to gene order rearrangements (Rocha, 2003). Reconfiguration of the ancestral angiosperm plastome through repeat-mediated recombination has now been reported in several groups (Sloan et al., 2014; Weng et al., 2014; Schwarz et al., 2015; Ruhlman et al., 2017; Choi et al., 2019; Choi et al., 2020). The recombinogenic potential of long repeats identified in the *Juncus* plastomes was likely involved in diversifying gene order.

The observation of slight variations in IR length between *Nicotiana* species was explored in seminal work that focused on the IR/LSC boundary in closely related groups. This work ultimately inferred recombination-mediated gene conversion between poly-A tracts that gave rise to a >12 kb expansion at the *N. acuminata* J_LB_ (IR_B_/LSC boundary) placing the new J_LB_ near *clpP* and duplicating the 12 kb sequence now included in the IR (Goulding et al., 1996). Although the details of the mechanism have been clarified and refined over the years, repeat-mediated gene conversion appears to be at the heart of it (Maréchal & Brisson, 2010; Oldenburg & Bendich, 2015; Ruhlman & Jansen, 2021).

Plastomes that contain a large number of long repeats can experience extensive rearrangement of gene order and both loss and gain of plastome sequence, including genes, introns and non-coding sequences alike. Expansion and contraction at both LSC and SSC boundaries contributed to variation in *Juncus* plastome size. Photosynthetic seed plant plastomes and IRs range from ~120-170 kb and 20-30 kb, respectively, however most IR-containing angiosperms sequenced to date display highly similar gene arrangement and plastome size (~150 kb; IR, ~25kb; Ruhlman & Jansen, 2021). Total plastome size in some groups is strongly influenced by IR expansion, yet in other lineages the association is loose at best. For example, a study of five *Cyperus* plastomes revealed the largest plastomes had the smaller IRs (i.e. *C. esculentus;* 186,255 kb/37,438 kb) and the smaller plastomes contained the larger IRs (i.e. *C. difformis;* 167,974 kb/38,427 kb) (Ren et al., 2021).

While total plastome size scaled with IR size (Table 2) and total repeat content (Table 3) in the four *Juncus*, the myriad events that altered each plastome relative to a shared ancestor with more conserved structure remain elusive. The smallest of the four plastomes, in *J. validus* would seem like a typical plastome based on the overall plastome and IR size (~147 kb and ~29 kb). However, the assembly and annotation show that it is not always size that matters. This plastome has likely experienced/is experiecing an ongoing series of IR boundary migrations resulting in a novel organization relative to the other taxa evaluated here. The near total elimination of the NDH gene suite, predominantly situated in the SSC in typical angiosperm plastomes, was unique to *J. validus* and suggests that IR boundary migration into the SSC played a role it their eventual loss. Although retained by the three other taxa, NDH sequences appear in alternate loci and several have been duplicated by IR inclusion (Figure 2; Figure S3) suggesting migration at the SSC boundaries. Indeed, the gene order arrangement proximal to IR/LSC boundaries display little rearrangement across all four *Juncus* (Figures 2, 3; Figure S3).

Complete ablation of the plastid-encoded NDH (NADH dehydrogenase-like) gene suite was reported for several unrelated seed plant lineages (Ruhlman et al., 2015). The NDH complex of plant and algal plastids participates in cyclic electron flow (CEF) (Shikanai et al., 1998) and comprises a multisubunit, plastid-localized complex that incorporates imported nuclear-encoded factors. The plastid genes encoding the NDH complex are highly conserved across Streptophyta (Hori et al., 2014) suggesting an essential function in photosynthesis (Ifuku et al., 2011). Using plastome sequencing and nuclear transcriptomics revealed that taxa lacking the plastid genes encoding constituents of NDH concomitantly lacked the relevant nuclear-encoded factors. Probing nuclear transcriptomes revealed that regardless of the state of the plastid NDH gene suite, genes encoding the alternate PGR5-dependent CEF pathway (Shikanai, 2014) were present in the nucleus of all examined taxa (Ruhlman et al., 2015). The loss of the NDH suite from the *J. validus* plastome is unique among examined Poales plastomes and suggest that an active PGR5-dependent pathway accounts for CEF in this species.

Apart from the loss of NDH genes, gene losses were shared by all four *Juncus* examined and included other genes that were lost from plastomes of diverse lineages (Ruhlman & Jansen, 2018). The plastid-localized Acetyl-coenzyme A carboxylase (ACCase; prokaryotic) is another multisubunit protein complex that incorporates nuclear-encoded polypeptides and participates in fatty acid metabolism (Ohlrogge & Browse, 1995). The plastid *accD* encodes one subunit of the four-unit complex and was lost in numerous taxa, often those that experienced other gene loss and pseudogenization events (Ruhlman & Jansen, 2018). Because plastid ACCase activity was thought an essential function (Kode et al., 2005) *accD* loss in several groups suggested that it may be expressed from a functional transfer to the nucleus or substituted by a redundant, nuclear-encoded enzyme (Konishi et al., 1996). In *Trifolium*, which lacks plastid *accD*, a functional transfer to the nucleus was uncovered (Magee et al., 2010). Further investigation failed to detect any remnant of the *accD* sequence in the plastomes of *T. repens* or *T. pratense* while mutated copies were identified in *T. aureum* and *T. grandiflorum* (Sabir et al., 2014). The 15-amino acid C-terminal catalytic domain of the ACCD protein, which is minimally required for prokaryotic ACCase function (Lee et al., 2004), was identified in the mutated copies and may indicate functionality. Probing nuclear transcriptomes from *T. repens* and *T. pratense* revealed that, as in *T. subterraneum* (Magee et al., 2010), a putatively functional ACCD protein was being expressed from a fusion sequence that included the ACCD catalytic domain (~270 aa) fused to the plastid target peptide from nuclear-encoded, plastid-targeted LPD1 (493 aa). Probing transcriptomes of related legumes that contained intact plastid *accD* was able to detect high identity copies of the ACCD core sequence suggesting that incorporation at nuclear loci predated the degradation of plastid *accD* (Sabir et al., 2014). Functional redundancy was demonstrated for prokaryotic ACCase (Babiychuk et al., 2011; Rousseau-Gueutin et al., 2013) and other gene products through transfer or substitution in different lineages (Ueda et al., 2007, 2008).

The fate of *accD* sequences and both the prokaryotic and the single-polypeptide eukaryotic ACCase in Poales has been a matter of investigation for some time. Morton & Clegg (1993) identified a recombination hotspot in seven Poaceae plastomes in the region between *rbcL* and *psaI* (i.e. the locus containing *accD* sequences in non-Poaceae plastomes (Harris et al., 2013). Exploiting the fact that both the eukaryotic and prokaryotic ACCases contain biotinylated polypeptides, Konishi et al. (1996) were able to identify which form of the enzyme was active in plastids from across the diversity of the green plant lineage, including two non-photosynthetic representatives. Differentiating the two enzymes by molecular weight revealed that only one group examined did not contain the 35 kDa peptide that represented the prokaryotic holoenzyme: Poaceae. Closer examination of Poales using PCR product sequencing combined with Southern blots probed with plastid *accD* from Commelinaceae taxa demonstrated pseudogenization or deletion in representatives of three families, Restionaceae, Joinvilleaceae and Poaceae (Harris et al., 2013). Extending the loss of *accD* to include the Cyperaceae (Cyperus, Ren et al., 2021; Elocharis, Lee et al., 2020) and now Juncaceae suggests either extreme lability of the coding sequence in Poales or that this gene was transferred or substituted by a nuclear encoded activity in a common ancestor. Differential nuclear retention, expression and transport of the gene product back to plastids among the various lineages could result in relaxed selection on the plastid gene (Ueda et al., 2007; Park et al., 2017).

The opportunity to sample deeply across and within lineages is revealing that the unusual variation identified by early Southern blots and more recent plastome sequencing suggests that these ‘unusual’ structural changes are not unique. The suite of plastid genes that are susceptible to pseudogenization or loss appears consistent across photosynthetic seed plants. Understanding phylogeny, inherent to evolutionary studies, requires deep sampling, high-quality sequencing, assembly and alignment to infer relationships. As next generation sequencing and single-molecule long-read sequencing platforms expand and become more accessible, reads will be generated for many diverse taxa. Where long sequence repeats exceed insert sizes in next gen systems, long reads will be able to ‘bridge the gap’. The ability to translate raw sequence reads into usable data for evolutionary and functional inquiries depends on advanced computational tools that provide fast, flexible platforms without vast computational demand. Facilitating this effort, the ptGAUL pipeline provides a fast and easy tool for assembling plastomes from long-read data, which will enable the characterization of repeat-rich, highly rearranged plastomes.

## ACKNOWLEDGEMENTS

We thank the UNC greenhouse staff for maintaining living *Juncus* materials and the UNC herbarium staff for storing our voucher specimens. We are very grateful for the UNC longleaf high-performance cluster for computational resources. This work was supported by National Science Foundation IOS-2034929 to A.M.J and C.D.J.

## DATA ACCESSIBILITY AND BENEFIT-SHARING

Demultiplexed sequence data of short-read and long-read data are available for download from the NCBI Sequence Read Archive (SRA) (BioProject PRJNA865266). The accession numbers of *J. roemerianus* and *J. validus* are OP235509 and OP235510, respectively. Information related to ptGAUL can be fetched in GitHub (https://github.com/Bean061/ptgaul).

## AUTHOR CONTRIBUTIONS

WZ developed the ptGAUL pipeline assembled the *Juncus* plastome and prepared most of the first draft of the manuscript. CEAF assembled downloaded, publicly available reads using ptGAUL and made modification to the script. Chaehee provided *Eleocharis dulcis* long-read data and confirmed the analyses on the rearrangement events of plastome and helped annotate the *Juncus* plastomes. Ruisen Lu helped analyze the long repeats and SSR numbers in *Juncus* and polished the annotation result for NCBI. Jeremy Wang helped modified the ptGAUL script. Robert Jansen and Tracey Ruhlman helped discuss and write the introduction and discussion on plastome rearrangement events. Alan Jones and Corbin Jones are the senior corresponding authors guiding this project and they polished the prose.

