## supplementary figures for "Plastid Genome Assembly Using Long-read Data (ptGAUL)"

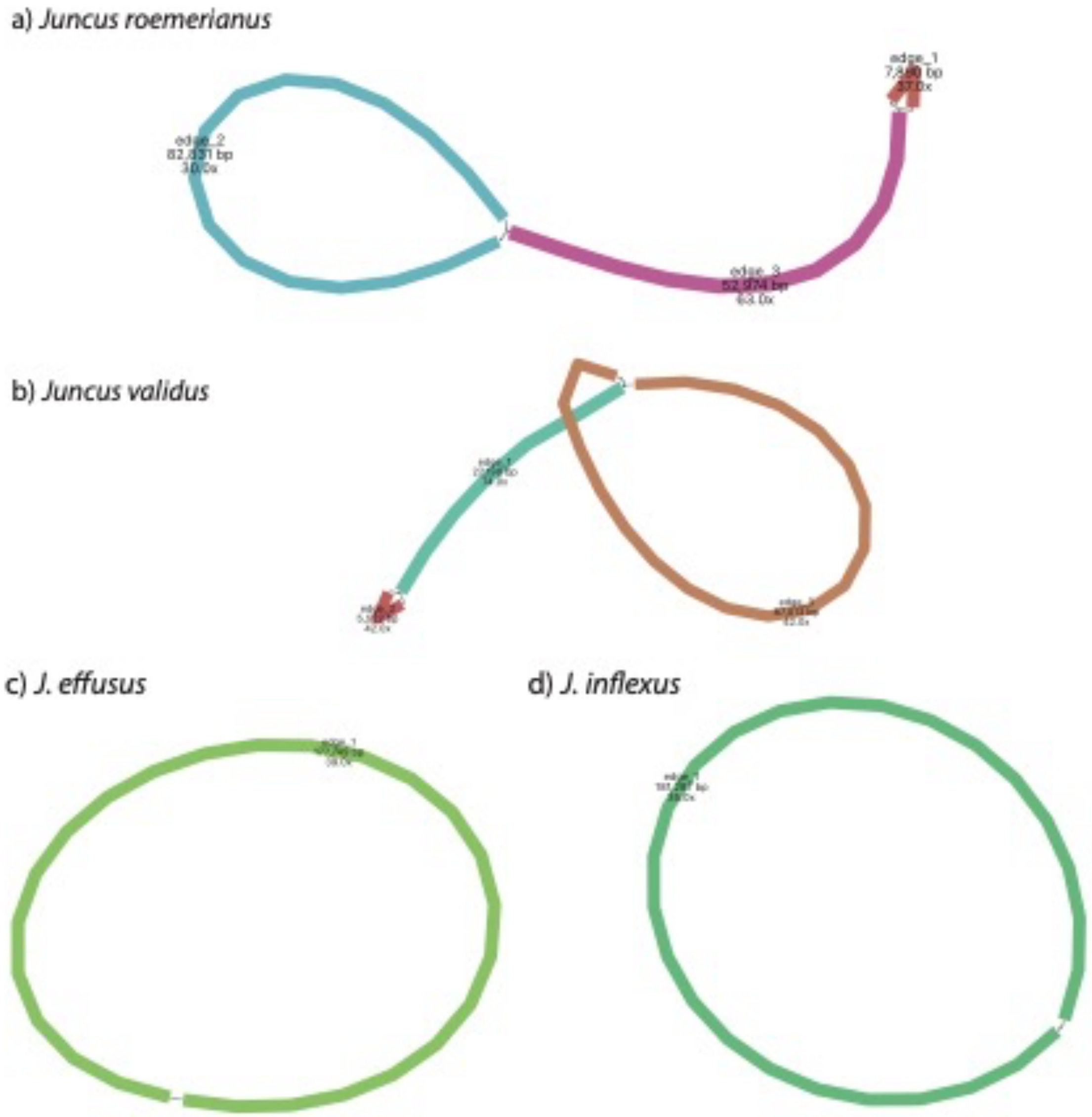


**Figure S1** plastome long-read assembly result of four *Juncus* from our study visualized by Bandage. Both a) *Juncus roemerianus* and b) *Juncus validus* have three assembled contigs; while c) *J. effusus* and d) *J. inflexus* form into circular contig. These circles and lassos of c) and d) represents major pattern of plastid chromosomes.

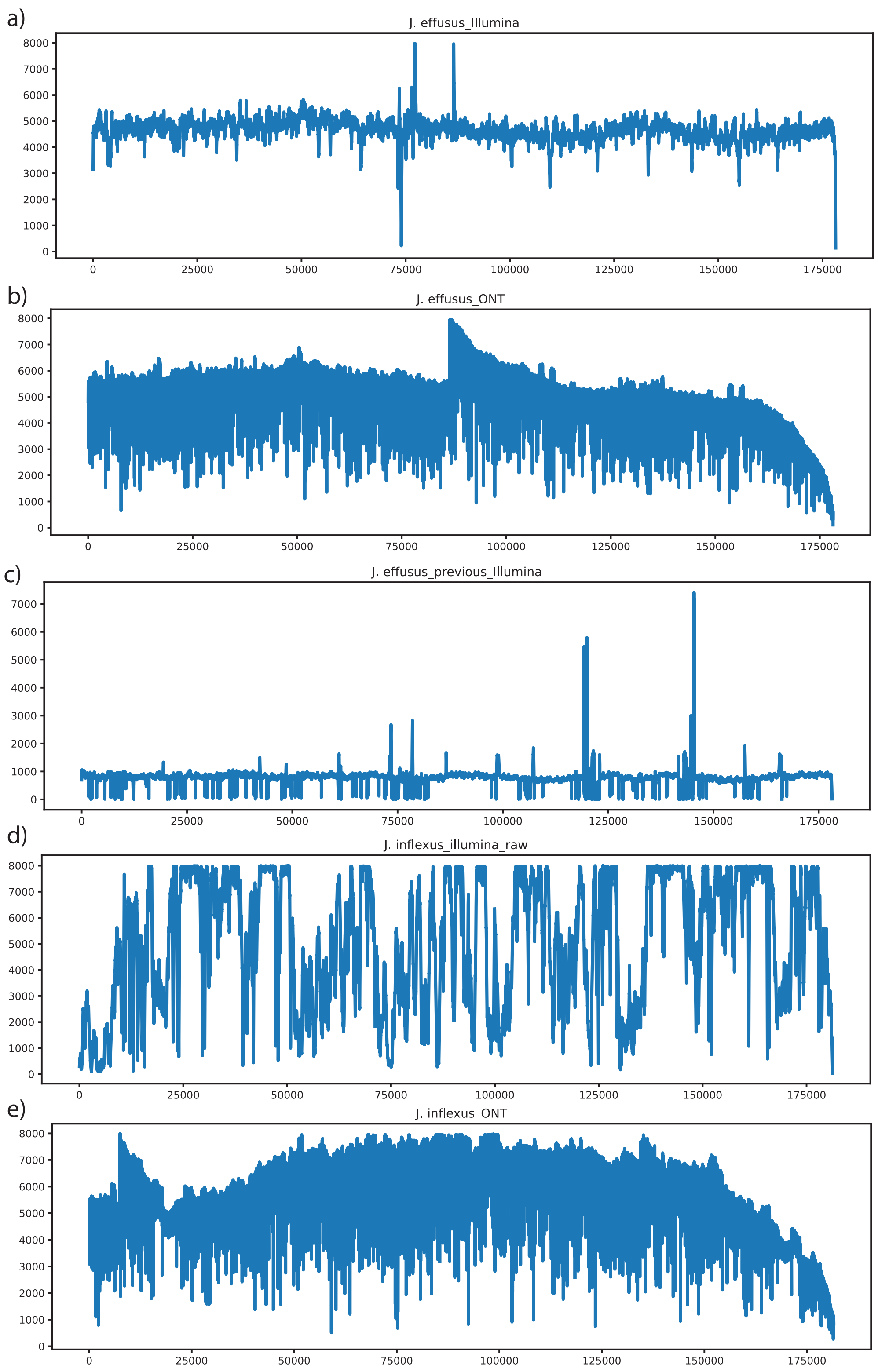


**Figure S2** Reads mapping result of two *Juncus* from minimap2 and samtools. A) Illumina short reads mapping result of *Juncus effusus* using the reference assembled from ptGAUL. B) ONT long reads mapping result of *Juncus effusus* using the reference assembled from ptGAUL. C) the mapping result of mapping Illumina read from Lu et al. (2021) to the reference assembled from ptGAUL using ONT reads from Planta et al. (2022) d) Illumina short reads mapping result of *Juncus inflexus* using the reference assembled from ptGAUL. e) ONT long reads mapping result of *Juncus inflexus* using the reference assembled from ptGAUL. *The Juncus inflexus* and *J. effusus* data of a), b), d), and e) were generated by Planta et al. (2022). The x axis represents the position of plastome, while the y axis means the coverage of each plastid position.

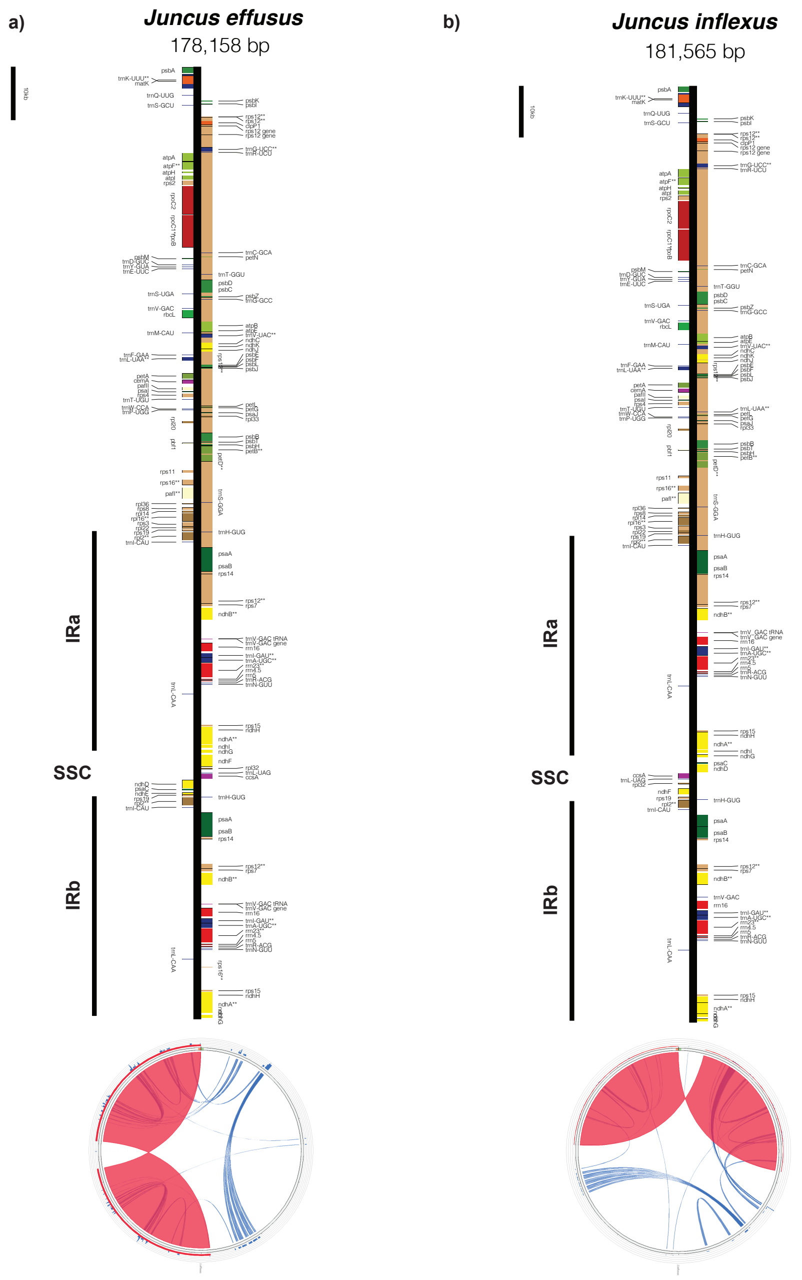


**Figure S3** Plastome structural maps of *J. effusus* and *J. inflexus*. a) and b) linear representations of *J. effusus* and *J. inflexus* plastomes, respectively, were drawn by OGDRAW. Genes that belong to different functional groups are color-coded. Small single copy (SSC), large single copy (LSC), and inverted repeats (IRa, IRb) are indicated for both plastid genomes. Circular maps of the two *Juncus* plastomes were used to evaluate repetitive DNA were depicted by Circoletto. The blue lines represent dispersed repeats in the plastome, while red regions represent the IR regions.

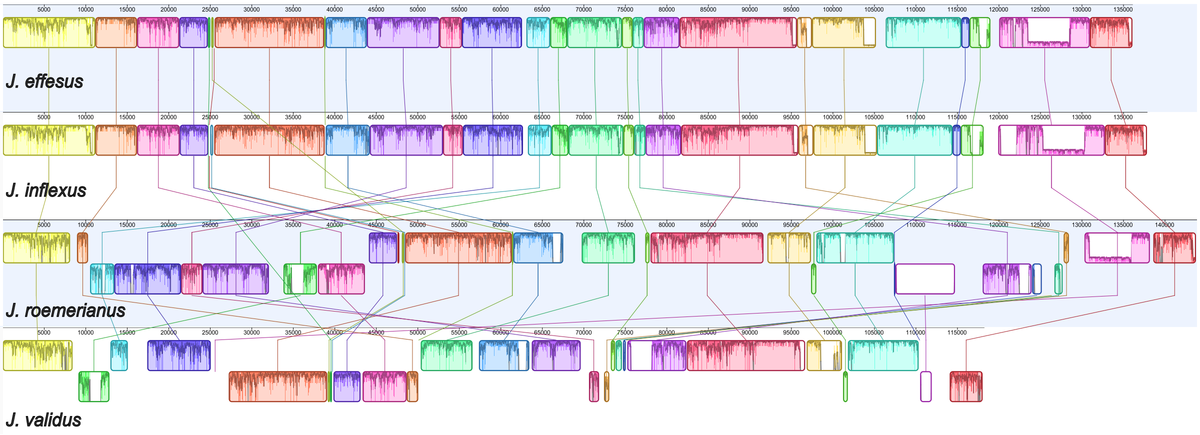


**Figure S4** Whole plastome alignment of four *Juncus* species. The LCBs were identified by progressiveMauve with *J. effusus* plastome as the reference. The corresponding LCBs among four plastomes are shaded and connected with a line of the same color. LCBs that are flipped over means inverted.

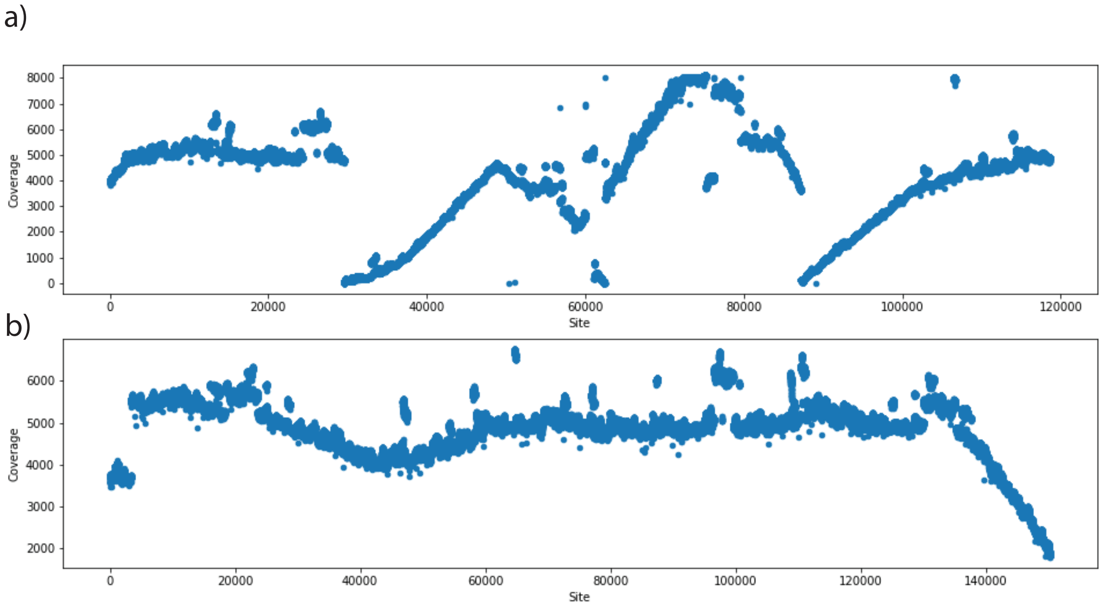


**Figure S5** Illumina reads mapping result of *Arctostaphylos glauca* from minimap2 and samtools. We A) mapped the Illumina reads to the plastome reference assembly by Huang et al. (2022). B) mapped the Illumina reads to the plastome reference assembly using ptGAUL in this study. The x axis represents the position of plastome, while the y axis means the coverage of each plastid position.

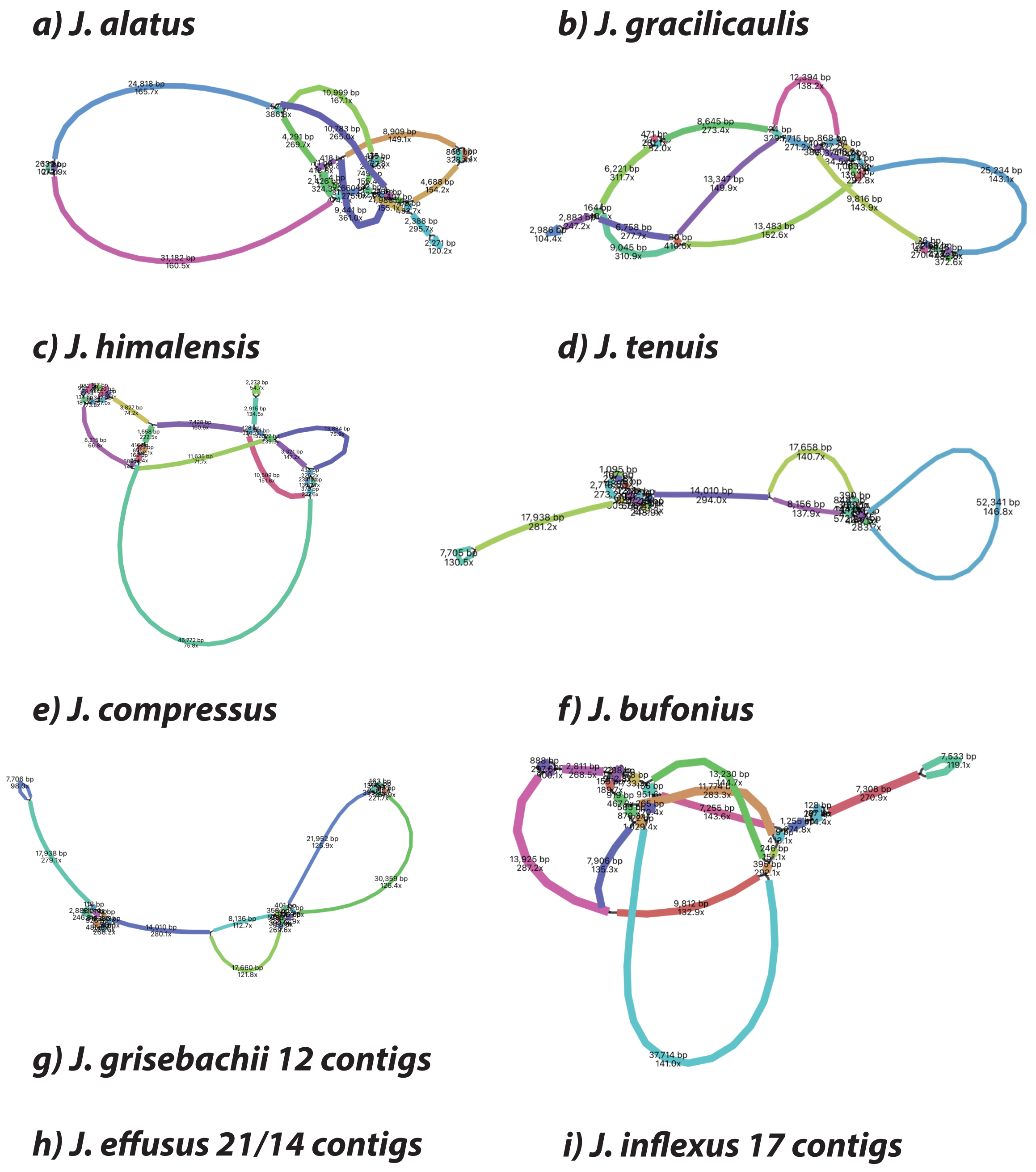


**Figure S7** plastome short-read assembly result of nine published *Juncus* species visualized by Bandage. Juncus species from a) to J) have numerous assembly paths detected; while species from g) to i) have fragmented contigs.
